# Sleuthing biochemical evidence to elucidate unassigned electron density in a CBL/SLAP2 crystal complex

**DOI:** 10.1101/2020.10.08.331397

**Authors:** Leanne E. Wybenga-Groot, C. Jane McGlade

## Abstract

The Src-like adaptor proteins (SLAP/SLAP2) bind to CBL E3 ubiquitin ligase to downregulate antigen, cytokine, and tyrosine kinase receptor signaling. In contrast to phospho-tyrosine dependent binding of CBL substrates through its tyrosine kinase binding domain (TKBD), CBL TKBD associates with the C-terminal tail of SLAP2 in a phospho-independent manner. To understand the distinct nature of this interaction, a purification protocol for SLAP2 in complex with CBL TKBD was established and the complex crystallized. However, determination of the complex crystal structure was hindered by apparent SLAP2 degradation during the crystallization process, such that only CBL TKBD residues could be modeled initially. Close examination of the CBL TKBD structure revealed a unique dimer interface that included two short segments of electron density of unknown origin. To elucidate which residues of SLAP2 to model in this unassigned density, a co-expression system was generated to test SLAP2 deletion mutants and define the minimal SLAP2 binding region. As well, SLAP2 degradation products were analyzed by mass spectrometry. Model building and map generation features of the Phenix software package were employed, leading to successful modeling of the C-terminal tail of SLAP2 in the unassigned electron density segments.

## 1. Introduction

The amino-terminal tyrosine kinase binding domain (TKBD) of CBL E3 ubiquitin ligase consists of a four-helix bundle (4H), an EF-hand, and a variant SH2 domain. The variant SH2 binds CBL substrates, such as phosphorylated tyrosine kinases, in a canonical phospho-tyrosine dependent manner(Meng *et al*., 1999). CBL TKBD also binds Src-like adaptor proteins (SLAP and SLAP2) and this interaction is important for CBL regulation of antigen, cytokine, and tyrosine kinase (TK) signaling (Sosinowski *et al*., 2000, Loreto & McGlade, 2003, Loreto *et al*., 2002, Sosinowski *et al*., 2001, Myers *et al*., 2006, Dragone *et al*., 2006, Dragone *et al*., 2009, Naramura *et al*., 1998, Lebigot *et al*., 2003, Pakuts *et al*., 2007). SLAP2 is composed of adjacent SH3 and SH2 domains most closely related to those found in SRC family kinase HCK, followed by a carboxy (C)-terminal tail region void of apparent domains or protein-protein interaction motifs and predicted to be predominantly disordered (Pandey *et al*., 1995, Tang *et al*., 1999, Wybenga-Groot & McGlade, 2013, 2015, Loreto *et al*., 2002). While the SLAP2 SH3/SH2 domains mediate binding to activated receptor TKs, the C-terminal tail region of SLAP2 associates with CBL TKBD in a phospho-tyrosine independent manner(Loreto & McGlade, 2003, Holland *et al*., 2001, Myers *et al*., 2006), suggesting that the interaction between CBL and SLAP2 is distinct and likely represents a novel mode of binding. Indeed, SLAP2 binding to CBL is not disrupted by a CBL Gly306Glu mutation, which abolishes TKBD binding to tyrosine phosphorylated substrates (Loreto *et al*., 2002). To understand the molecular basis of this interaction, we purified and crystallized CBL TKBD in complex with SLAP2. Initial efforts to determine the structure of the CBL/SLAP2 complex resulted in a structure of a CBL dimer, with two unassigned regions of electron density nestled in the dimer interface that resembled alpha helices. This paper describes how we used biochemical and mass spectrometry techniques to determine which residues of SLAP2 to model in this unknown, unassigned electron density.

## 2. Materials and methods

### 2.1 Cloning, expression and purification

Residues 28-259 of mouse SLAP2 or 29-261 of human SLAP2 were cloned in frame into a modified pET32a (pET32a-mod) vector (with a TEV cleavage site downstream of the His tag and 17 residues upstream of the thrombin cleavage site; gift from Gil Privé lab, UHN) with BamHI/BglII and XhoI sites, to express thioredoxin (Trx)-His(6)-mSLAP2 and (Trx)-His(6)-hSLAP2. Residues 25-357 of CBL were cloned in frame into pGEX4T-1 to express glutathione-S-transferase (GST)-CBL. Using standard QuikChange methods, 19 residues downstream of the TEV cleavage site were deleted from Trx-His(6)-hSLAP2, thus abolishing the thrombin cleavage site and placing the TEV cleavage site in closer proximity to the protein N-terminus, to generate Trx-His(6)-Δlinker-hSLAP2. Trx-His(6)-hSLAP2 or Trx-His(6)-Δlinker-hSLAP2 were cloned in frame into multiple cloning site (MCS) 1 of pETDuet-1 (Novagen), with CBL (47-357) cloned in frame into MCS2 of the same pETDuet-1 vector, for co-expression of SLAP2 and CBL (Duet-His-hSLAP2-CBL and Duet-His-Δlinker-hSLAP2-CBL (261)). Standard QuikChange site-directed mutagenesis methods were employed to delete residues 255-261 from SLAP2 in Duet-His-Δlinker-hSLAP2-CBL (254) and delete residues 198-229 and 255-261 from Duet-His-hSLAP2-CBL (29-261 Δ198-229 and 29-254 Δ198-229).

Trx-His(6)-mSLAP2 was transformed into *Escherichia coli* BL-21 cells and grown in 8 L Luria Bertani (LB) media supplemented with 50 μg/mL ampicillin (Amp) overnight at 16°C (A_600_= 0.6-0.9 and 0.5 mM isopropyl-β-D-thiogalactopyranoside (IPTG) induction). Cells were collected by centrifugation, cell pellet frozen at −80°C, thawed, resuspended in 150 mL lysis buffer (50 mM HEPES pH 7.5, 0.5 M NaCl, 10% glycerol, 10 mM imidazole, 10 mM β-mercaptoethanol (βME), 10 mM MgCl_2_, 5 mM CaCl_2_, cOmplete EDTA-free protease inhibitor cocktail tablets (inhibitor tablets) (Roche Applied Science), benzonase nuclease, 1 mM phenylmethylsulfonyl fluoride (PMSF)) and lysed by three cycles of high pressure homogenization (Emulsiflex) and two cycles of sonication on ice. Following centrifugation, supernatant was mixed with Ni-NTA agarose (Qiagen) for 90 min by gentle nutating at 4°C. Resin was washed with 6 x 25 mL of wash buffer (50 mM HEPES pH 7.5, 0.5 M NaCl, 10% glycerol, 20 mM imidazole, 10 mM βME, 10 mM MgCl_2_, 5 mM CaCl_2_) and SLAP2 protein eluted with wash buffer containing increasing concentrations of imidazole (75, 150, 225, 300 mM imidazole, 28 mL total elution volume). The Trx-His(6) tag was cleaved by addition of 300 units of thrombin (Sigma T4648) directly to the eluate and the solution dialyzed (Slide-A-Lyzer dialysis cassettes, Thermo Scientific, 3500 MWCO) overnight at room temperature (rt) against 2 L dialysis buffer (25 mM HEPES pH 7.5, 0.4 M NaCl, 4% glycerol, 10 mM imidazole, 5 mM βME, 5 mM MgCl_2_, 5 mM CaCl_2_). The solution was removed from the dialysis cassette, centrifuged at 4000 rpm for 7 min to remove precipitate, and PMSF added to the soluble portion to 1 mM. This SLAP2 solution was passed very slowly over the same aliquot of Ni-NTA resin (washed since elution) to remove Trx-His(6), concentrated with a centrifugal filter unit (Amicon Ultra, Millipore), and flash frozen in liquid nitrogen for storage at −80°C.

To express CBL TKBD protein, GST-CBL was overexpressed and cells harvested as above (except 6.5 hrs at 37°C, 1 mM IPTG). Cell pellet was resuspended in 100 mL lysis buffer (50 mM HEPES pH 7.5, 0.5 M NaCl, 10% glycerol, 10 mM βME, 2 mM MgSO_4_, inhibitor tablets, benzonase nuclease) and lysed as above. Following centrifugation, supernatant was mixed with Glutathione Sepharose 4B (GE Healthcare) for 90 min by gentle nutating at 4°C. Resin was washed with 6 x 25 mL of wash buffer (50 mM HEPES pH 7.5, 0.5 M NaCl, 10% glycerol, 10 mM βME, 5 mM CaCl_2_) and 150 units of thrombin (Sigma #T4648) added directly to the resin and left overnight at 4°C; thrombin addition and incubation was repeated at rt. CBL protein was eluted by washing resin with elution buffer (25 mM HEPES pH 7.5, 0.4 M NaCl, 5% glycerol, 5 mM βME), concentrated as above, and stored stably at 4°C for weeks.

Approximately 33 mg of CBL or 10 mg of mSLAP2 were loaded individually onto a HiLoad 26/60 Superdex 75 gel filtration column (GE Healthcare Life Sciences) equilibrated with GF buffer (25 mM HEPES pH 7.5, 0.4 M NaCl, 2% glycerol, 5 mM βME) and protein elution detected by monitoring A_280_. Fractions corresponding to isolated CBL or SLAP2 protein were assessed for purity by 12.5% SDS-PAGE analysis. To isolate the CBL/SLAP2 complex, purified CBL and SLAP2 protein were mixed in a 1:1 molar ratio, concentrated to 1 mL, and loaded onto the same gel filtration column as above. Fractions corresponding to co-elution of CBL and SLAP2 were assessed for purity by 12.5% SDS-PAGE analysis and concentrated to 11.2 mg/mL. (Concentration was estimated by measuring the absorbance of the protein at 280 nm and calculating its concentration using a combined CBL/SLAP2 molecular extinction coefficient of 94810 M^−1^ cm^−1^ as predicted by ProtParam (http://web.expasy.org; (Gasteiger *et al*., 2003))).

For co-expression of CBL and SLAP2/SLAP, Duet-His-Δlinker-hSLAP2-CBL constructs 261 and 254 and truncation constructs Duet-His-hSLAP2-CBL 29-261 Δ198-229 and 29-254 Δ198-229 were overexpressed in *E. coli* BL-21 (100 mL LB, 50 μg/ml Amp, overnight growth at 16-18°C, and 0.35-0.5 mM IPTG induction). Cell pellets were resuspended in SLAP2 lysis buffer as above, except 25 mM HEPES pH 7.5, 150 mM NaCl, lysed by sonification, and purified as above on Ni-NTA agarose (Qiagen) with wash buffer 25 mM HEPES pH 7.5, 150 mM NaCl, 2% glycerol, 20 mM imidazole, 10 mM βME. Protein was eluted as above with increasing imidazole concentrations or bead slurry was mixed with SDS 2X loading buffer (125 mM Tris pH 6.8, 4% SDS, 20% glycerol, 0.7 M βME, bromophenol blue), boiled, and analyzed by 12.5% SDS-PAGE, stained with Coomassie blue. Eluted protein was dialyzed, cleaved, Ni-NTA clarified, and concentrated as above for SLAP2 (except 25 mM Hepes pH 7.5, 0.4 M NaCl, 5% glycerol, 10 mM βME, 5 mM CaCl_2_). The CBL/SLAP2 complex was further purified by gel filtration chromatography and assessed for purity as above, with GF buffer (except 5% glycerol).

### 2.2 Crystallization of the CBL/SLAP2 complex

The co-eluted CBL/SLAP2 complex was mixed at 5 mg/mL in a sitting drop with equal volume (200 nL) of 0.1 M Hepes pH 7.5, 10% (w/v) PEG8000 at rt using a Mosquito robot (TTP Labtech). Small rod-like crystals were observed after approximately five months. A solution of 50% glycerol was added to the drop prior to harvesting and flash freezing the crystals in liquid nitrogen.

### 2.3 Data collection and processing for CBL/SLAP2 complex

Diffraction data was collected at the Advanced Photon Source and processed with Mosflm and QuickScale (Pointless, Aimless/Scala, Ctruncate) software to 2.9 Å(Winn *et al*., 2011, Evans, 2011). Initial attempts at molecular replacement (MR) were performed with Phenix software (Phaser_MR) (Adams *et al*., 2010) using CBL TKBD (PDB id: 2CBL), one SH3 domain from Lck (PDB id: 1LCK), and one SH2 domain from Lck (PDB id: 1LCK) as a model. The same diffraction data was re-processed as above to 2.5 Å and subsequent attempts at MR performed with Phaser_MR and CBL TKBD (PDB id: 1B47) as a model. Anisotropic scaling of the data was performed (services.mbi.ucla.edu/anisoscale) followed by multiple iterative cycles of model building in Coot and refinement with Phenix_refine (Emsley & Cowtan, 2004, Adams *et al*., 2010, Emsley *et al*., 2010). A Feature Enhanced Map (FEM) (Afonine *et al*., 2015) was calculated using Phenix software and poly-glycine chains placed in the unassigned density using the “Find Helices and Strands” feature in Phenix, the new FEM, and the improved model. Poly-glycines were manually converted to poly-alanines, and then to mSLAP2 residues ^240^GLRESLSSYISLAEDP^255^ in both chains of unassigned density. Real space refinement was performed with Coot and refinement with Phenix. Using the FEM, residues ^237^LSE^239^ were assigned. Symmetry related molecules and superpositions were calculated in Coot.

### 2.4 Mass spectrometry

SDS-PAGE gel bands were reduced, alkylated, digested with trypsin protease as per SPARC BioCentre’s in-gel digestion protocol (https://lab.research.sickkids.ca/sparc-molecular-analysis/services/mass-spectrometry/mass-spectrometry-sample-protocols/). Digested peptides were subjected to LC-MS/MS at SPARC BioCentre (The Hospital for Sick Children) (60 min gradient, Thermo LTQ Orbitrap) and raw data searched with Mascot and X!Tandem software against the human proteome, with carbamidomethylation (C) as a fixed modification and pyro-glu (Q, N-terminal E), s-carbamoylmethylcysteine cyclization (N-terminus), deamidation (NQ), oxidation (M) and acetylation (N-terminus) as variable modifications.

### 2.5 Isothermal titration calorimetry

Purified CBL TKBD protein and peptides representing mSLAP2 C-terminal tail (LSSYISLAEDPLD), phosphorylated kinase activation loop of Lyn kinase (VIEDNEpYTAR), and phosphorylated Zap70 kinase (TLNSDGpYTPEPA) were employed in isothermal titration calorimetry (ITC) experiments, performed on a VP-ITC MicroCalorimeter (MicroCal, Northampton, MA). CBL TKBD protein and peptide solutions were made in 20 mM Hepes pH 7.0, 200 mM NaCl and degassed just prior to titrations. The reference cell was filled with degassed distilled water. Aliquots of 10 μL of SLAP2 or Lyn-pY peptide at 137-155 μM were injected at 3-4 min intervals into the sample cell, which contained CBL at 14 μM. Aliquots of 10 μL of Zap70-pY peptide at 223 μM were injected at 5 min intervals with CBL at 21 μM in the sample cell. Titration curves were analyzed using the Origin software provided by MicroCal.

## 3. Results and Discussion

To understand the molecular basis of SLAP2 adaptor protein binding to CBL TKBD, mouse SLAP2 protein (residues 28-259) and CBL TKBD protein (residues 25-357) were purified by affinity and size exclusion chromatography (Fig. 1A,B). To form a complex of CBL TKBD and SLAP2 for co-crystallization studies, the purified proteins were mixed in a 1:1 molar ratio and subjected to size exclusion chromatography (Fig. 1C). The proteins eluted as two peaks, one at an earlier volume than typically observed for either CBL TKBD or SLAP2, indicating higher molecular weight (Fig. 1C). As confirmed by SDS-PAGE of chromatography fractions, the first peak contained a CBL/SLAP2 complex (Fig. 1D), which was concentrated and used for sparse matrix crystallization experiments. Small rod-like crystals (Fig. 1E) diffracted to 2.5 Å resolution, albeit with strong anisotrophy, at the Advanced Photon Source. After integrating and scaling the diffraction data (Table 1), the Matthews coefficient formula predicted that the asymmetric unit contained one CBL TKBD and one SLAP2 molecule. However, when Phaser_MR searched for a solution, with one molecule of CBL TKBD (PDB id: 2CBL), one SH3 domain from Lck (PDB id: 1LCK), and one SH2 domain from Lck (PDB id: 1LCK) (the structure of the SLAP2 SH3/SH2 module had not yet been solved), the translation function Z (TFZ) score and log-likelihood gain (LLG) were low (TFZ 4.2-9.9), indicating a poor molecular replacement (MR) solution. In contrast, when Phaser_MR searched for a solution with two molecules of CBL TKBD, the TFZ score increased to 15.7, suggesting a correct solution had been found. Limited refinement of the model resulted in a R-factor of 0.375 and Rfree of 0.424. Examination of crystal packing and symmetry related molecules suggested insufficient open space for full-length SLAP2 (Fig. 2AB). Given the structure of CBL TKBD had been solved and published numerous times already, further refinement of this model was not immediately pursued. However, superposition of the Cα atoms of one molecule from our CBL TKBD model with one CBL TKBD molecule from structures available in the protein data bank at the time (eg. PDB id 3BUX, 3BUW, 3BUM) revealed a similar mode of configuration or packing of CBL molecules in the published structures, and a distinct mode of interaction in our model. In the published structures, interactions between CBL molecules are mediated through the EF-hands (Fig. 2C), while the dimer interface in our model involves the EF-hand and the 4H bundle (Fig. 2D). More importantly, unassigned electron density resembling two alpha helices appeared in the space between the two CBL TKBD molecules in our model (Fig. 2EF). Although the quality of the density and its helical sidechains was not sufficient to establish sequence registry, we reasoned that the density likely represented a portion of the SLAP2 protein that co-crystallized with CBL. However, detailed understanding of SLAP2 structure and mapping of its interaction with CBL required to model SLAP2 residues into the density were not available.

**Table 1.**
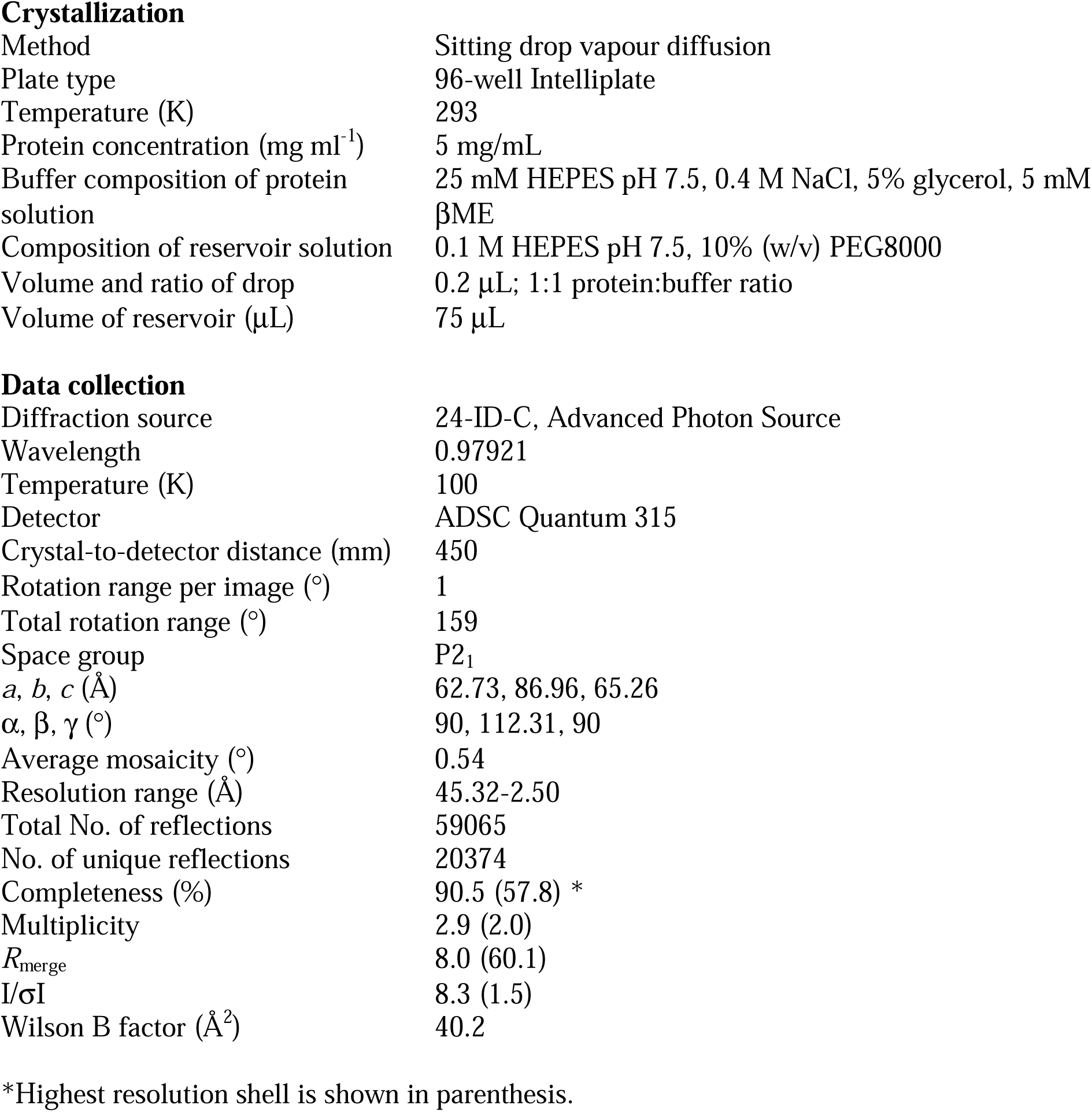
Crystallization, x-ray diffraction and data collection statistics.

|  |  |
| --- | --- |
| Method | Sitting drop vapour diffusion |
| Plate type | 96-well Intelliplate |
| Temperature (K) | 293 |
| Protein concentration (mg ml<sup>-1</sup>) | 5 mg/mL |
| Buffer composition of protein solution | 25 mM HEPES pH 7.5, 0.4 M NaCl, 5% glycerol, 5 mM βME |
| Composition of reservoir solution | 0.1 M HEPES pH 7.5, 10% (w/v) PEG8000 |
| Volume and ratio of drop | 0.2 μL; 1:1 protein:buffer ratio |
| Volume of reservoir (μL) | 75 μL |
| **Data collection** |  |
| Diffraction source | 24-ID-C, Advanced Photon Source |
| Wavelength | 0.97921 |
| Temperature (K) | 100 |
| Detector | ADSC Quantum 315 |
| Crystal-to-detector distance (mm) | 450 |
| Rotation range per image (°) | 1 |
| Total rotation range (°) | 159 |
| Space group | P2<sub>1</sub> |
| *a*, *b*, *c* (Å) | 62.73, 86.96, 65.26 |
| $\alpha$, $\beta$, $\gamma$ (°) | 90, 112.31, 90 |
| Average mosaicity (°) | 0.54 |
| Resolution range (Å) | 45.32-2.50 |
| Total No. of reflections | 59065 |
| No. of unique reflections | 20374 |
| Completeness (%) | 90.5 (57.8) * |
| Multiplicity | 2.9 (2.0) |
| *R*<sub>merge</sub> | 8.0 (60.1) |
| *I*/σ*I* | 8.3 (1.5) |
| Wilson B factor (Å<sup>2</sup>) | 40.2 |

\*Highest resolution shell is shown in parenthesis.

**Figure 1:**
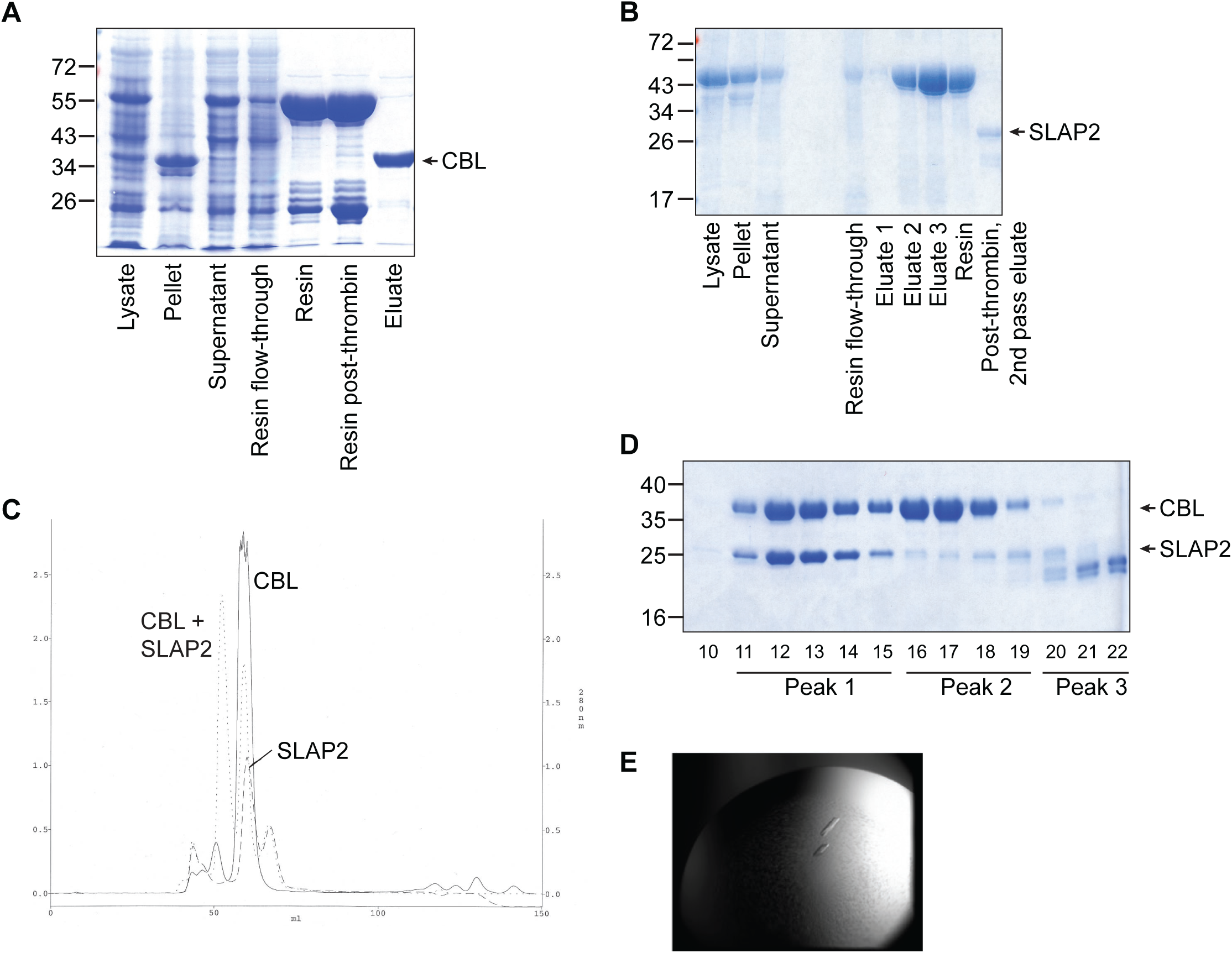
Purification and crystallization of CBL TKBD and SLAP2. A) SDS-PAGE gel stained with Coomassie blue of samples from stages of CBL TKBD protein purification. B) As in A, for mouse SLAP2. C) Absorbance chromatograms (280 nm) for CBL TKBD (solid line) and mSLAP2 (hatched line) proteins subjected to gel filtration analysis individually and mixed in a 1:1 molar ratio (dotted line). D) SDS-PAGE gel stained with Coomassie blue of fractions 10-22 from gel filtration analysis of the CBL/SLAP2 complex. E) Crystals formed from CBL/SLAP2 complex.

**Figure 2:**
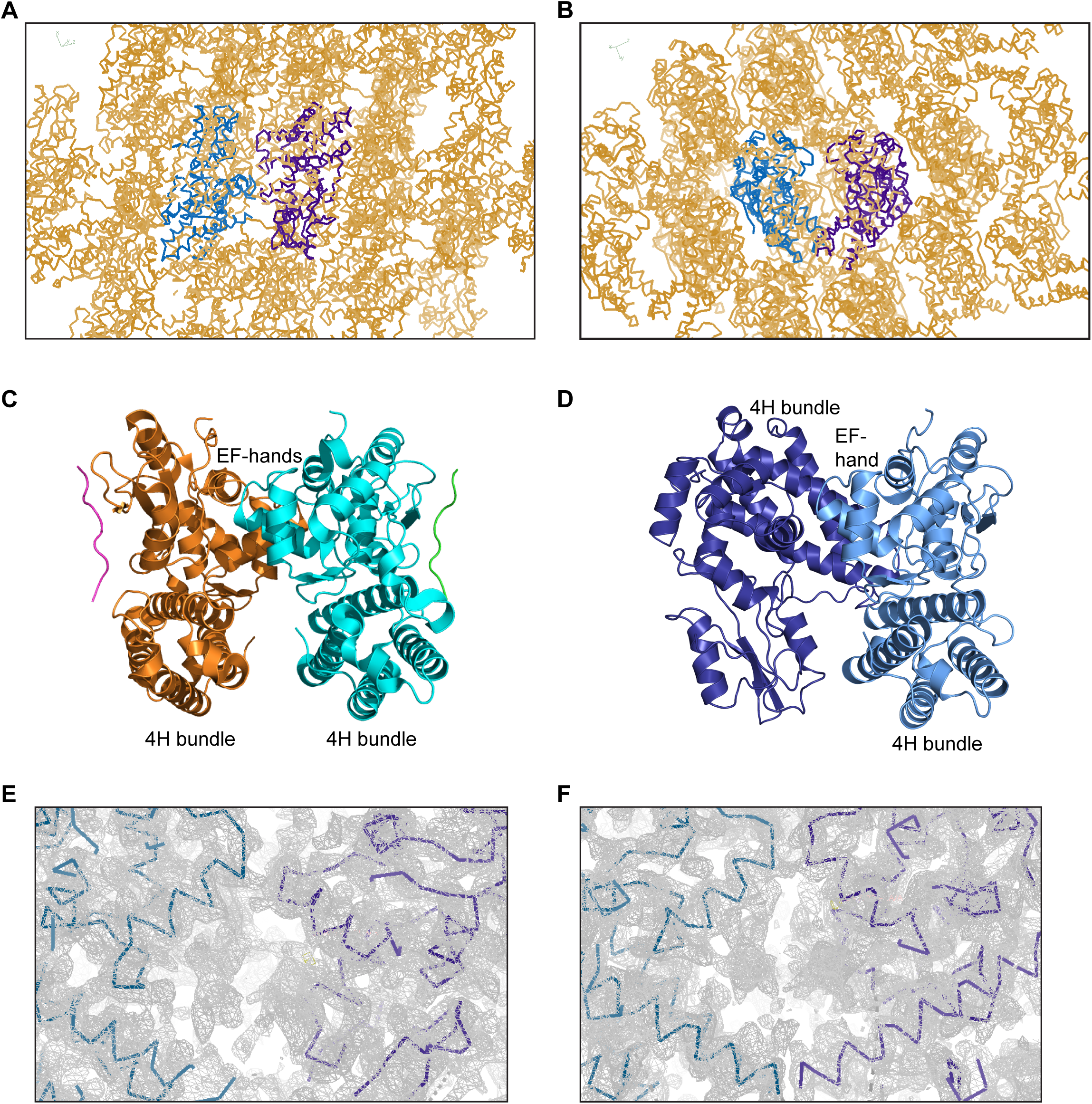
Distinct crystal packing of CBL TKBD monomers. A) Cα traces of the CBL TKBD crystal structure, with the Cα trace of CBL molecules 1 and 2 shown as light and dark blue lines and symmetry related molecules as calculated by Coot shown in orange. B) As in A, different orientation of the dimer interface. C) Ribbon representation of the Cα atoms of a representative CBL structure (PDB id:3BUW), with molecules 1 and 2 shown in cyan and orange, respectively, pTyr peptides shown in green and magenta, and the TKBD 4H bundles labelled. D) Ribbon representation of the Cα atoms of the CBL TKBD crystal structure, with molecules 1 and 2 shown in light and dark blue, respectively, and the TKBD 4H bundles labelled. Molecule 1 is in approximately the same orientation as that in C. E) Electron density at the CBL dimer interface is shown in grey, with the Cα trace of CBL molecules 1 and 2 shown as light and dark blue lines. F) As in E, rotated approximately 90° about the horizontal axis.

To simplify the purification of SLAP2 in complex with CBL TKBD and further define the SLAP2 binding region, we generated a Duet co-expression vector system. The human SLAP2 sequence (residues 29-261) was preceded by thioredoxin (Trx) and His-tag fusion proteins to allow co-purification of CBL TKBD that was bound to Trx-His-hSLAP2 during the initial Ni-NTA affinity chromatography purification step (Fig. 3A). Following digestion with thrombin to remove fusion proteins and subsequent elution from the Ni-NTA resin, the CBL/SLAP2 complex was further purified by gel filtration chromatography (Fig. 3B). While additional crystallization trials with this pure CBL/SLAP2 complex were unsuccessful, we observed that fractions that eluted at a higher volume and no longer contained CBL protein also appeared to contain degradation products of SLAP2. This suggested that the degradation products could no longer bind CBL TKBD, especially since they were not present in the fractions where CBL and SLAP2 co-eluted. To determine the sequence of the degradation products, we performed in-gel protease digestion on the corresponding gel bands (Fig. 3B, red boxes) and analyzed the peptides by liquid chromatography tandem mass spectrometry (LC-MS/MS). Notably, many peptides from the N-terminal portion of the SLAP2 C-tail were identified (Fig. 3C, grey box), while no peptides from the C-terminus were identified in either product (Fig. 3C, white box). The peptides identified from the two degradation products differed only at the N-terminus of SLAP2. These data suggest that SLAP2 degraded at its C-terminal end during the purification process and that this degraded protein could no longer bind CBL TKBD. These observations identified hSLAP2 residues 243-261 as important for CBL binding.

**Figure 3:**
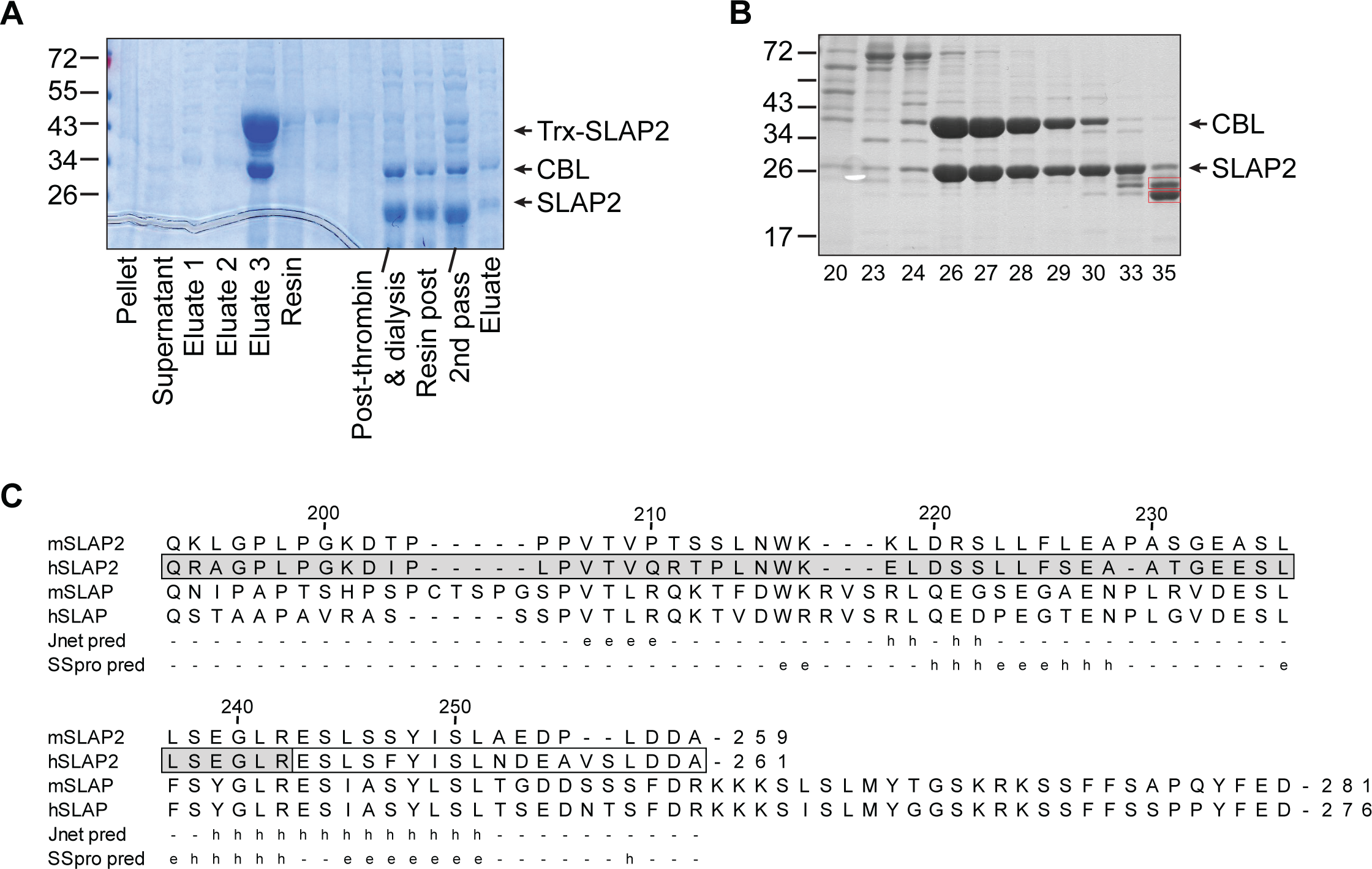
Co-purification of CBL/SLAP2 and mass analysis of degradation products. A) SDS-PAGE gel stained with Coomassie blue of samples from stages of CBL/SLAP2 protein co-purification. B) As in A, for fractions 20-35 from gel filtration analysis of the CBL/SLAP2 complex. Red boxes indicate bands excised for in-gel digestion and LC-MS/MS analysis. C) Sequence alignment of the C-tail region of SLAP/SLAP2 proteins, with residue numbering based on human SLAP2. Secondary structure predictions for mSLAP2 as calculated by Jnet (Jnet pred) and Scratch (SSpro pred) are shown, with e, -, and h representing β-strand, coil and α-helix, respectively. Grey box represents residues identified by LC-MS/MS analysis. White box represents residues not found in LC-MS/MS experiments.

With this knowledge in hand, we purchased a synthetic 13-mer peptide representing the C-terminal tail of mSLAP2 (^245^LSSYISLAEDPLD^257^) for binding studies. Isothermal titration calorimetry (ITC) experiments failed to show association between purified CBL TKBD and the SLAP2 peptide (Fig. 4A). In contrast, phospho-tyrosine peptides from Zap70 and Lyn kinases, which bind the variant SH2 domain of CBL TKBD, bound purified CBL TKBD in ITC experiments performed under similar conditions (Fig. 4BC). This indicated that the 13-mer SLAP2 peptide is not sufficient for CBL binding.

**Figure 4:**
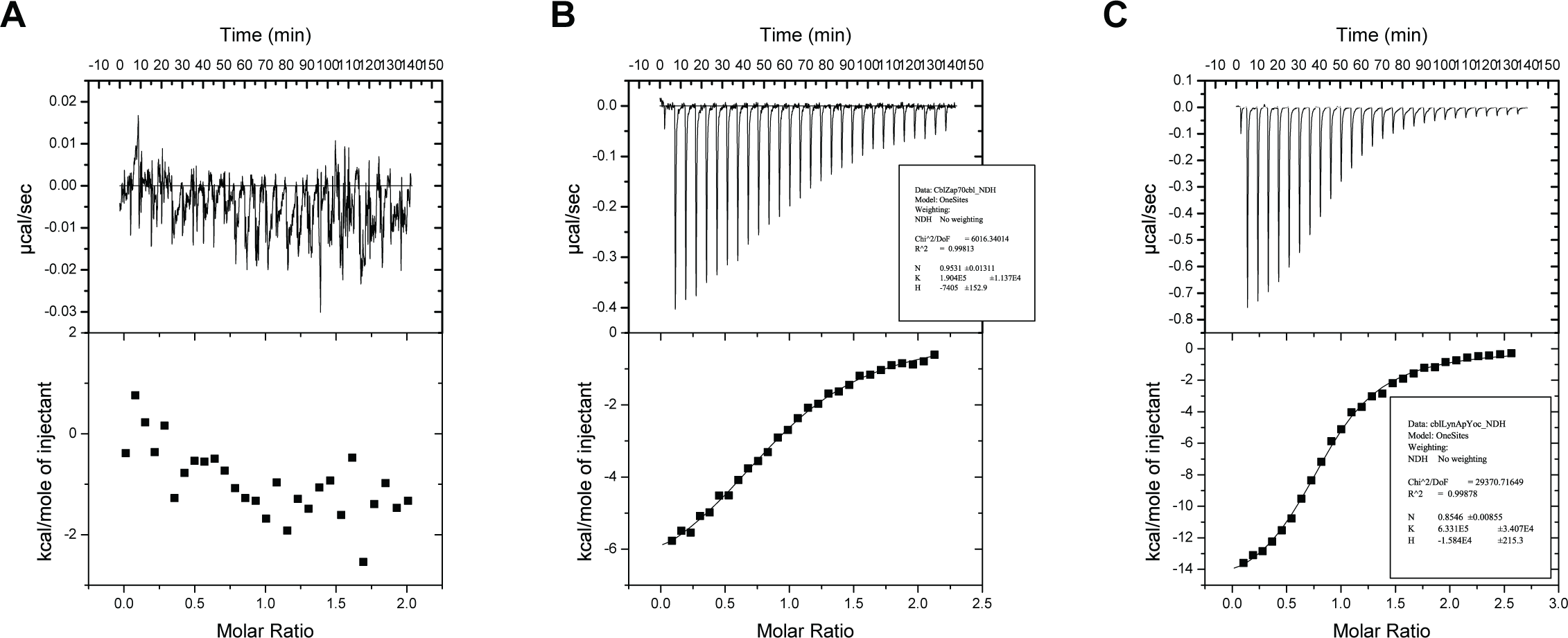
Isothermal titration calorimetry with CBL TKBD and peptides. ITC data for CBL TKBD with A) mSLAP2 peptide, B) Zap70 pY peptide, and C) Lyn pY peptide. The top panels show the heat evolved upon injection of peptide into CBL protein, plotted as a function of time. Peaks correspond to individual injections. Bottom panels show normalized integrated heats of reaction plotted against the molar ratio of total ligand concentration to total protein concentration. The solid line represents the best fit to the data, according to a one-site binding model.

Large portions of the SLAP2 C-terminal tail are expected to be disordered by secondary structure and disorder prediction programs (Fig. 3C) (Drozdetskiy *et al*., 2015, Cheng *et al*., 2005). Knowing that disordered regions can hinder crystallization and structure determination, we sought to better understand the minimal SLAP2 region required for interaction with CBL. To this end, we generated truncation constructs in our Duet co-expression vector and tested their ability to co-purify CBL TKBD. Residues 198-229 and/or 255-261 were deleted from the human SLAP2 C-terminal tail based on sequence conservation and predictions. Deletion of these C-terminal regions did not disrupt co-purification of CBL TKBD (Fig. 5). This defined the region that is critical for CBL binding to residues 230-254 of hSLAP2. This is consistent with mass spectrometry analysis of the SLAP2 degradation products and secondary structure prediction programs which predict either alpha helical structure or a combination of alpha helical and β-strand structure in this region (Fig. 3C).

**Figure 5:**
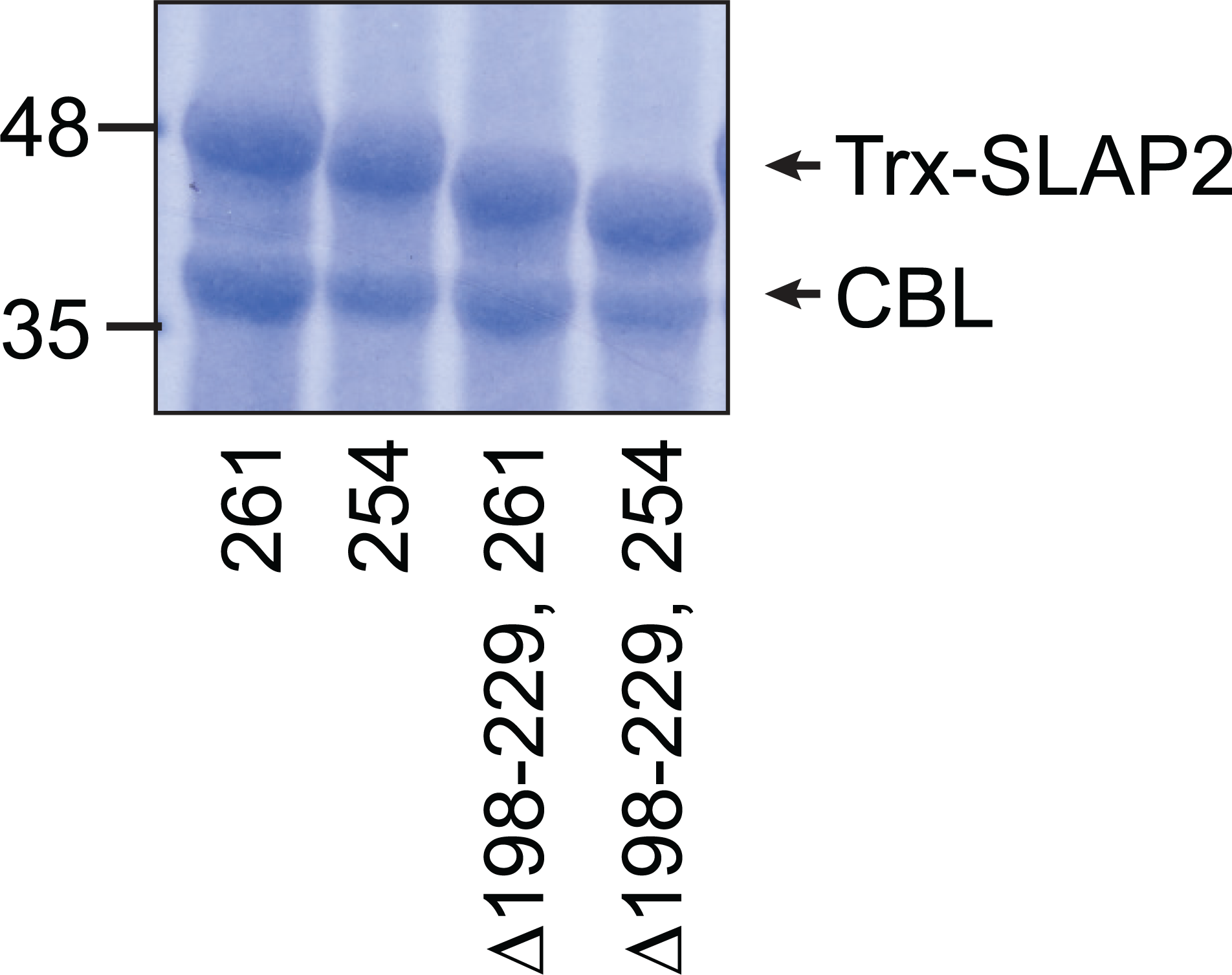
Co-purification of CBL/SLAP2 truncation constructs. SDS-PAGE gel stained with Coomassie blue for CBL and SLAP2 proteins co-purified from Duet-His-Δlinker-hSLAP2-CBL (261), its truncated version (254), Duet-His-hSLAP2-CBL 29-261 Δ198-229, and its truncated version 29-254 Δ198-229.

Having better defined which residues of SLAP2 are both necessary for CBL binding and predicted to have secondary structure, we returned to the original CBL/SLAP2 co-crystallization x-ray diffraction data and model. First, the data was integrated and scaled again, this time at 2.5 Å based on processing statistics (Table 1) and limits determined by anisotropic scaling (2.5, 2.8, 2.5 Å) (Strong *et al*., 2006). Our initial attempts at structure determination by molecular replacement employed CBL 47-351 in complex with a Zap70 phospho-peptide (PDB id: 2CBL). However, phospho-peptide binding to the CBL SH2 domain is known to induce a closure of the domains, with a slight shift and rotation of the SH2 domain toward the 4H bundle (Meng *et al*., 1999). Indeed, superposition of the Cα atoms of the SH2 domain of our CBL TKBD model with that of native (PDB id: 1B47) versus liganded TKBD (PDB id: 2CBL) showed greater similarity to unliganded TKBD. A subsequent attempt at MR with native CBL TKBD (PDB id: 1B47) as a model produced TFZ and LLG scores of 19.2 and 522.2, respectively, indicative of an improved solution. After multiple iterative cycles of refinement with Phenix.refine, and model building in Coot and Phenix with the “Phase and build” feature(Emsley & Cowtan, 2004, Adams *et al*., 2010), the Rfree was 33.0%. A Feature Enhanced Map (FEM)(Afonine *et al*., 2015) was calculated using Phenix software, which further improved the quality and interpretability of the electron density map (Fig. 6AB). Poly-glycine chains were placed in the unassigned density using the “Find Helices and Strands” feature in Phenix, the new FEM, and the improved model. Poly-glycines were manually converted to poly-alanines and then to mSLAP2 residues ^240^GLRESLSSYISLAEDP^255^ in both chains of unassigned density based on the contour of the density (Fig. 6C). Real space refinement in Coot fit the SLAP2 residue side chains within the previously unassigned density, and refinement in Phenix reduced the Rfree to 31.2%. Using the FEM, residues ^237^LSE^239^ were manually modeled in the last fragment of unassigned density.

**Figure 6:**
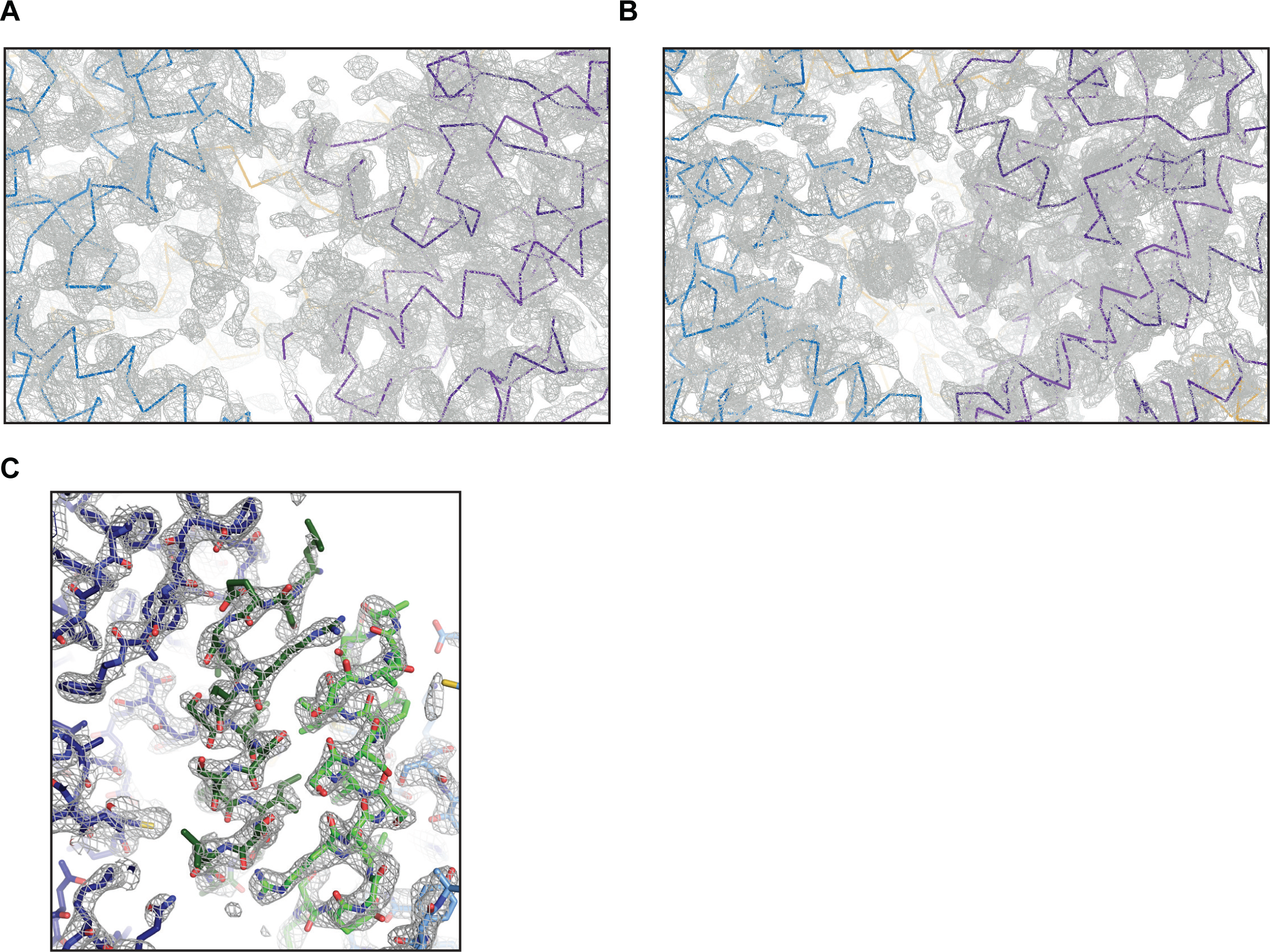
Modeling SLAP2 residues in unassigned, α-helical electron density. A) Electron density at the CBL dimer interface is shown in grey, with the Cα traces of CBL molecules 1 and 2 shown as light and dark blue lines, respectively. B) As in A, rotated approximately 90° about the horizontal axis. C) Electron density at the site of CBL and SLAP2 interaction is shown in grey, with CBL and SLAP2 atoms shown in stick. Carbon atoms are coloured according to their respective backbones, with CBL monomers in blue and SLAP2 monomers in green. Oxygen and nitrogen atoms are coloured red and blue, respectively. For clarity, portions of CBL in the plane of the page have been removed.

It was not possible to establish if additional SLAP2 residues co-crystallized with CBL TKBD but remained disordered. However, based on protein packing within the crystal (Fig. 2AB) and our observation that SLAP2 is susceptible to degradation during the purification process, we conjecture that CBL/SLAP2 co-crystallized following degradation of the C-terminal tail from the SH3/SH2 domain. Based on our biochemical evidence and x-ray crystallographic data, we propose that residues 237-255 represent a critical component for CBL/SLAP2 interaction. It remains to be determined if the SH3/SH2 module and/or additional residues of the SLAP2 C-terminal region also associate directly with CBL TKBD.

## Acknowledgements

The authors thank Savar Kaul, Julia Manalil, Alissa Visram, and Mackenzie Kinney for technical assistance, Elton Zeqiraj for diffraction data collection, and Frank Sicheri for access to crystallization resources and diffraction data collection. This work was supported with funds from the Cancer Research Society, Garron Family Cancer Centre, and the Canadian Institutes of Health Research FRN166034(to CJM). LWG was support by fellowships from CIHR and American Brain Tumor Association.

## Author Contributions

LEW-G performed experiments and wrote the manuscript; CJM supervised LEW-G and wrote the manuscript.

